# DoOR 2.0 - Comprehensive Mapping of *Drosophila melanogaster* Odorant Responses

**DOI:** 10.1101/027920

**Authors:** Daniel Münch, C. Giovanni Galizia

## Abstract

Odors elicit complex patterns of activated olfactory sensory neurons. Knowing the complete olfactome, i.e. responses in all sensory neurons for all odorants, is desirable to understand olfactory coding. The DoOR project combines all available *Drosophila* odorant response data into a single consensus response matrix. Since its first release many studies were published: receptors were deorphanized and several response profiles were expanded. In this study, we add to the odor-response profiles for four odorant receptors (Or10a, Or42b, Or47b, Or56a). We deorphanize Or69a, showing a broad response spectrum with the best ligands including 3-hydroxyhexanoate, alpha-terpineol, 3-octanol and linalool. We include these datasets into DoOR, and provide a comprehensive update of both code and data. The DoOR project has a web interface for quick queries (http://neuro.uni.kn/DoOR), and a downloadable, open source toolbox written in R, including all processed and original datasets. DoOR now gives reliable odorant-responses for nearly all *Drosophila* olfactory responding units, listing 693 odorants, for a total of 7381 data points.

*At the time of uploading this preprint, a preview of the DoOR 2.0 webpage is available at:* *http://neuro.uni.kn/DoOR/2.0*

## Introduction

The coding capacity of even small olfactory sensory systems is enormous because of the multidimensional nature of the olfactory code (often referred to as combinatorial). The vast majority of odorants elicit specific response patterns across olfactory sensory neurons (OSN), consisting of strongly activated, weakly activated, inhibited and non activated OSNs [1, 2, 3]. In a simplified binary system (the “combinatorial” case), 50 OSNs would have a theoretical capacity of 2^50^ odors. To understand the principles of this code one would like to know the response profiles of all OSNs of a system for as many odorants as possible. Deorphanizing odorant response profiles is laborious and it is being performed in labs across the world with different technical approaches. This yields heterogeneous sets of data: some from electrophysiological single sensillum recordings, some from calcium imaging, some from heterologous expression systems, to name but a few. The fruit fly *Drosophila melanogaster* is, to date, the species with most deorphanized and well described odor-response profiles. With the DoOR project we have created a way to combine all the existing, heterogeneous odorant response data into a single consensus response matrix [4]. When DoOR was established in 2010 the response profiles of several receptors were still missing and the number of available odorant responses per receptor ranged from 7 to 184 with a mean of 74.

While DoOR uses *Drosophila* as a model organism, it addresses olfactory coding in general: The complexity of sensory systems differs across species but most show striking similarities in their neurocomputational logic [5, 6]. For example, sensory neurons in the mammalian nose or on an insect antenna express a single or a few receptor genes, defining their tuning towards a specific set of odorants. As a first approximation, the axons of OSNs that express the same receptors converge onto glomeruli in the first olfactory input center of the brain (the olfactory bulb in mammals or the antennal lobes in insects) [7, 8, 9, 10]. In some cases one cell type expresses more than one receptor [11, 12]. Therefore, there is no clear and universally valid definition of “information channel”: receptor gene, receptor protein combination, glomerulus: whichever definition of “information channel” has to allow for flexibility. Nevertheless, it is across this set of OSNs (or “information channels”) with their distinct but overlapping tuning to different odorants, that brains extract the olfactory information about the environment. The number of these information channels differs between animals: humans have ~ 400 functional genes that code for potential receptors, dogs ~ 800 and mice ~ 1000 [13]. In honeybees, OSNs project to ~ 160 different glomeruli and in ants up to 630 glomeruli were found [14, 15]. The fruit fly *Drosophila* melanogaste*r* has about ~ 50 OSN classes [7, 8, 16, 17].

DoOR has been used by the community as a basis for further analysis both in physiology and in modeling [18, 19, 20, 21, 22, 23, 24]. The DoOR web page (http://neuro.uni.kn/DoOR) has become an established source of information for the community: there were ~ 140 unique web sessions per month from ~ 50 countries in 2014. However, since 2010 a lot of new responses to odorants were published. These data fill important holes in the DoOR project and significantly extend our knowledge about the *Drosophila* olfactome, and therefore the time is ripe to integrate them into DoOR.

Here, we present a major update of the DoOR project in information and in calculation (DoOR 2.0). Nevertheless, we maintain data and functions as two separate packages and we add new versions in a transparent way, keeping all previous versions available. We now have 78 responding units (“information channels”) and 693 odorants with the number of odorants responses per responding unit ranging from 11 to 497 with a mean of 100 odorants per responding unit.

## Results

### Different types of new data

With this update of the DoOR.data package we introduce three classes of new data: (1) Better precision raw data from datasets that were already included. When we published the first version of DoOR, not all raw data from the considered publications was available. In many cases we had to estimate quantitative responses from graphical plots. In the meantime we received the raw data for many of these studies from the authors and updated the existing DoOR datasets. This situation is improving: publishing data that underlies the plots of a paper in the supplements becomes a widespread practice, giving increasingly access to raw data in the community. (2) Raw data from new publications: Many new studies have been published since the first DoOR release. These studies added important new data to the *Drosophila* olfactome. Importantly, they deorphanized most of the remaining OR response profiles for which no ligand was known previously. (3) Data recorded by us that is first published along with this paper. Here we present the response profiles of five OSN classes, measured via *calcium* imaging. Our set of ~ 100 odorants, added a total of 529 new odorant responses to DoOR. One of these response profiles deorphanizes the Or69a sensory neurons, others contribute to existing profiles (Figure 2). See Table S1 for an overview of all studies that contributed to the DoOR project. We note, with respect to points 1 and 2, that many but not all colleagues were willing to share (published) odorant response data.

We added original data for the following receptors (Figure 2 and Table S5): For **Or10a** OSNs methyl salicylate, ethyl benzoate and butyric acid elicited the strongest responses in our hands. About one third of the tested odorants led to a reduction of calcium concentration (inhibition). In total our dataset added 52 odorants to the known response profile. Or10a is expressed in ab1D neurons, their axons innervate glomerulus DL1. The cells co-express the gustatory receptor Gr10a [8]. For **Or42b** our data added 76 substances. 3-hexanone, ethyl propionate and ethyl (S)(+)-3-hydroxybutyrate were the three strongest ligands in this dataset. The receptor is chirality selective: the stereo isomer ethyl (R)-(-)-3-hydroxybutyrate did not activate these neurons. Or42b is expressed in ab1A neurons, their axons innervate glomerulus DM1. **Or47b** responded mainly with inhibition, confirming previous reports [25, 26, 3]. We added 61 new odorant responses to DoOR and observed weak inhibitions for most substances. Benzaldehyde, furfural and acetic acid produced stronger inhibitions. Excitatory responses were weak as compared to other receptor cells. The strongest responses in our set were to (S)-carvone and propanoic acid, but these are not the best ligands. A stronger ligand, methyl laurate, has been recently discovered by Dweck *et al*. [27], the dataset is included in DoOR. Or47b is expressed in at4 neurons, their axons innervate glomerulus VA1lm. **Or56a** expressing ab4B OSNs responded best to geosmin, confirming the deorphanizing data by Stensmyr *et al*. [28]. In addition, stimulation with (1R) (-)-fenchone and alpha-ionone also evoked lower but reliable signals. We also observed inhibitory responses. 2,3-butanedione and acetic acid produced the strongest decreases in fluorescence. In total we added 82 responses to the response profile. Or56a is expressed in ab4B neurons, their axons innervate glomerulus DA2. The cells co-express the receptor Or33a [8]. For **Or69a** no ligands were known previously. We recorded responses to 106 odorants. Or69a OSNs responded particularly broad, showing activity towards most of the odorants in our set: the receptor kurtosis was -0.36. We found the strongest responses for ethyl 3-hydroxyhexanoate, alpha-terpineol, 3-octanol and linalool. Or69a is expressed in ab9 neurons, their axons innervate glomerulus D. Overall, across the five characterized receptors, we analyzed the response profiles with respect to chemical class (Figure 2). It is apparent from the figure that chemical class is not a good response predictor: all colors intermingle across the entire odor-response range. In other words, it is not useful to characterize individual receptors as “alcohol receptor”, or “ester receptor”.

### Deorphanized OSNs

With the new datasets included, DoOR response profiles are now existing for all OSNs except for Ir40a, and all antennal lobe glomeruli except VA7m have been assigned to a sensory neuron (Figure 1b and Table 1). All other glomeruli without cognate receptor gene in the previous 2010 version of DoOR have been deorphanized now. They are innervated by IR expressing OSNs housed in coeloconic sensilla or in the sacculus [12]. We updated the “OSN to glomerulus” mappings and the glomerulus nomenclature according to Silbering *et al.*[12] and the recently published *in-vivo* atlas of the *Drosophila* AL by Grabe *et al.*[17]. Ligands have been published for all IRs [12], and are included in DoOR.

**Figure 1:**
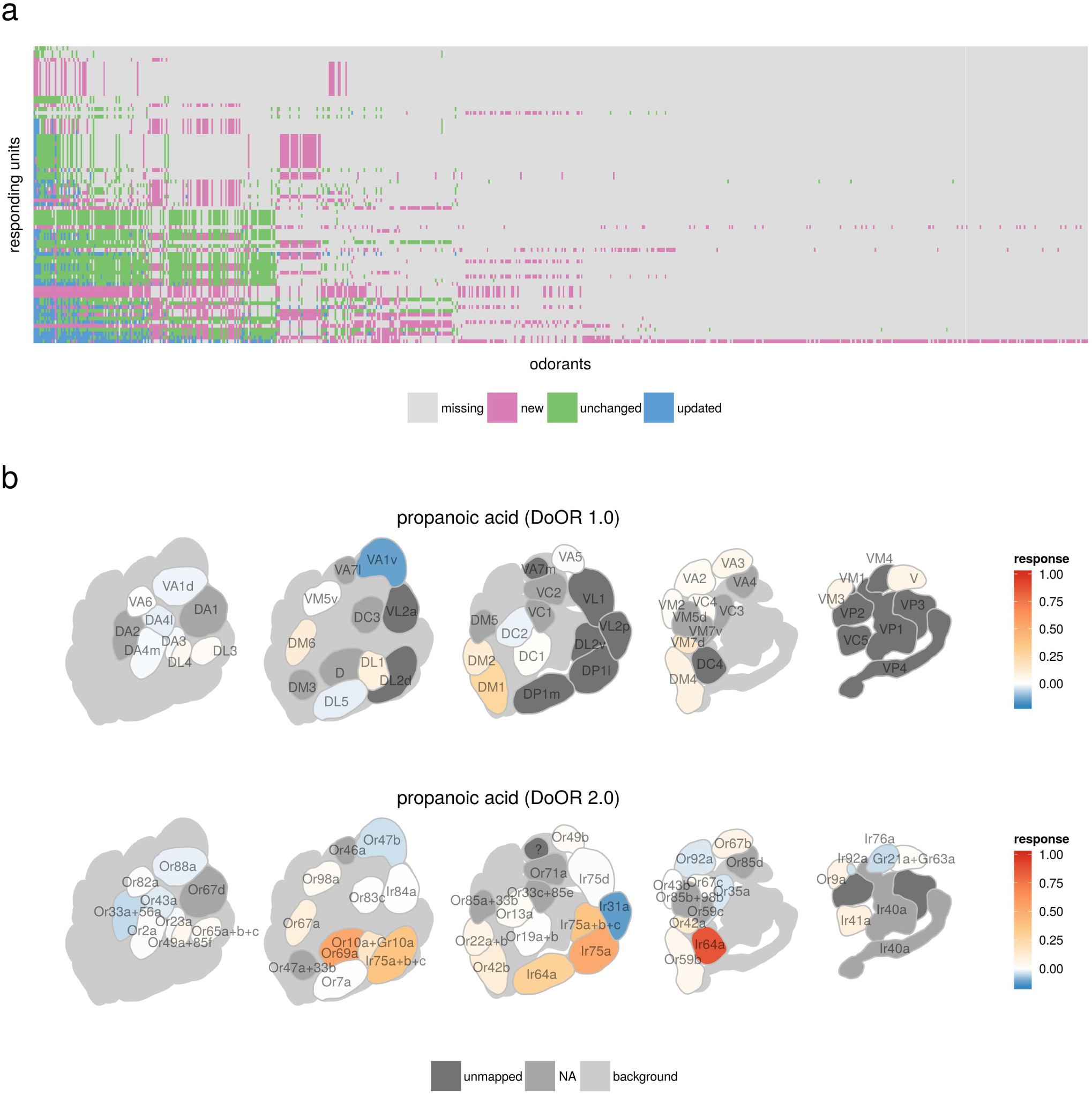
Changes in DoOR 2.0 as compared to the previous DoOR database. **a** Each point in the matrix represents an odor-responding unit combination. Colors indicate whether that combination is new in DoOR (*red*), was updated (*blue*), unchanged (*green*) or is still missing (*grey*). Response units were sorted according to the numbers of odorants they were tested with, odorants were sorted accordingly (see Tables S2, S3 and S4 for responding unit and odorant names respectively). **b** Visualization of ensemble responses. Responses elicited by propanoic acid mapped onto a representation of the *Drosophila* antennal lobe model from Grabe *et al.*[17]. Glomerulus names are shown in the *top* panel, the corresponding receptor names are shown in the *bottom* panel. DoOR 2.0 contains mappings of 13 IR innervated glomeruli that were still unmapped (*dark grey*) in DoOR 1.0. The two *dark grey* glomeruli VP2 and VP3 are non-olfactory.

**Figure 2:**
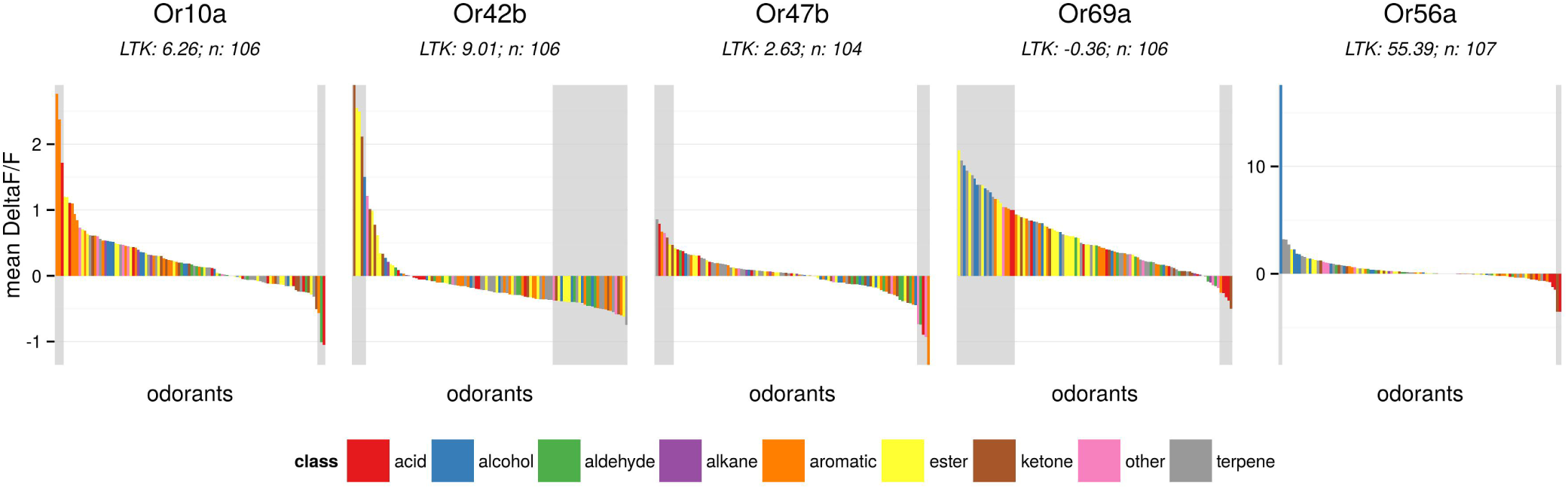
Response profiles for five OSNs measured *via* calcium imaging and added to DoOR. Bars represent mean calcium signals (n = 3-16 and 122-296 for controls) measured from five different GAL4 driver lines in response to a set of ~ 100 odorants. Colors indicate the chemical classes the different odorants belong to. Shaded areas indicate half maximal and half minimal response ranges respectively. Or56a was recorded using a different reporter (GCaMP3 vs. GCaMP1.3); the different scale is due to the reporter, and does not indicate different receptor calcium response properties. Mineral oil solvent responses were subtracted. Number of odorants in the dataset is given as *n*. All odorant names and response values are given in Table S5.

**Table 1:**
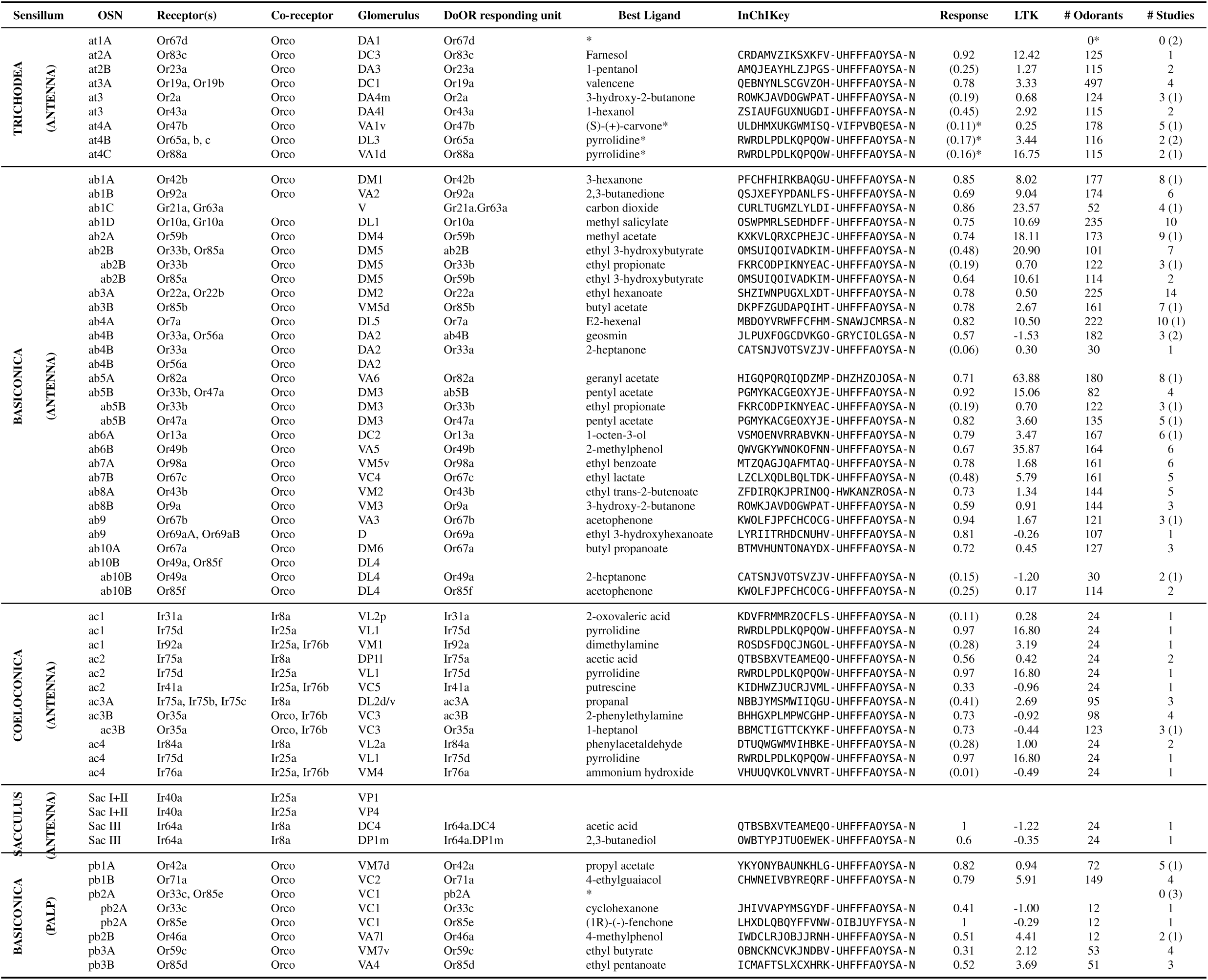
Overview of receptors, corresponding OSN classes, and targeted glomeruli,with the DoOR responding unit nomenclature. The best ligand in theDoOR database is given with its InChIKey. Sorted by sensillum/OSNname. *LTK,* lifetime kurtosis, see Methods for calculation;*Response,* the consensus response of the best ligand withthe SFR subtracted, weak responses (below 0.5) are shown in parentheses;*#Odorants*, number of odorants that are included in the final consensusresponse matrix; *#Studies*, numbers in parentheses indicatestudies that had to be excluded due to low merge quality or too lowoverlap with other studies; *\**, possible stronger ligandexisting in an excluded study; other responding units in DoOR: `ac1,ac1A, ac1B, ac1BC, ac2, ac2A, ac2B, ac2BC, ac3_noOr35a, ac4Or1a,Or22c, Or24a, Or30a, Or45a, Or45b, Or59a, Or74a, Or85c, Or94a, Or94b`.

### Updated DoOR mappings

There were multiple cases where response profiles could not be assigned unambiguously to a single receptor already in DoOR 1.0. These included cases where the receptor of the measured OSN was not known (e.g. OSN ac1A), or where more than one functional receptor was expressed in an OSN (e.g. ab5B, which expresses Or47a and Or33b). In these cases we assigned the response profile to the OSN name. While some of these cases have been resolved in the meantime (unknown partners were mostly IRs), others remain. Consequently, we had to expand this naming scheme, assigning some response profiles to the sensillum and others to the glomerulus they were measured in. Due to these difficulties in nomenclature we refer to the different origins of DoOR response data as “responding units” throughout the text. In most - but not all - cases a “responding unit” consists of an unambiguous mapping of receptor cell in a given sensillum, receptor gene/protein, and glomerulus in the antennal lobe. All “responding units” are listed in Table 1, together with the relevant information.

In many cases electrophysiological recordings from coeloconic sensilla could not be mapped to the individual OSN, because spike amplitudes and/or shape were not discrete enough to perform a separation. For example, single sensillum recording (SSR) data from Silbering *et al.*[12] was integrated as summed OSN responses for the individual coeloconic sensilla ac1-4. Marshall *et al.*[29] were able to separate the unit with the strongest amplitude (the A neuron) but summed the remaining. We considered this data as e.g. “ac1A” (the strongest, unambiguous) and “ac1BC” (the other two, not separable). Mapping responses recorded from glomeruli to the corresponding IR is also not always straight forward due to complex innervation patterns. For example, OSNs that express Ir64a project from chamber III in the sacculus to the two glomeruli DC4 and DP1m [30]. As a mapping to the IR name would be ambiguous and the OSN names for ac sensilla are not well defined, we extended our nomenclature and introduced concatenated names of the receptor & glomerulus (Ir64a.DC4 and Ir64a.DP1m). Ir75d is expressed in three different OSNs housed in the three sensilla ac1, ac2 and ac4, they all target the VL1 glomerulus. Assuming that the IR is the main determinant of the OSN response, we mapped recordings from the VL1 glomerulus to Ir75d. We also used the published IR response profiles to estimate putative sensillum/receptor/glomerulus relationships (see below).

We updated existing names in two instances. We renamed Gr21a to Gr21a.Gr63a as neither receptor alone is functional and none is known to be a co-receptor [31]. We renamed ab3B to Or85b as the co-expression of Or98b is not clear and no response profile for the latter is existing [7]. Or23a and Or83c as well as Or2a, Or19a and Or43a were initially described as being expressed in the trichoid sensilla at2 and at3 [7]. Ronderos *et al.*[32] describe Or23a and Or83c OSNs as being housed in an intermediate sensillum and thus rename at2 to ai2. Dweck *et al.*[33] renamed at2 and at3 to ai1 and ai2 respectively. Since these two proposed new nomenclatures are conflicting we decided to reduce confusion and to keep the old at1-at4 nomenclature.

### Merging algorithms

Apart from code optimization which increases computational speed, the core algorithms for merging several datasets into a single consensus response profile remained unchanged. In short: datasets were rescaled to the range [0,1] and merged pairwise by calculating the best fitting function on odorants recorded in both studies. Odorants measured only in one of the studies were subsequently projected onto this function. The sequence of merging was determined by iteratively finding the pairs that produced the best fit [4]. The “best fit” was quantified as the fit yielding the lowest “mean orthogonal distance” (MD, see Methods). Where feasible (*n_datasets_* ≦ 10 ≡ 3,628,800 permutations) we computed all possible merging sequences and selected the one with the lowest MD to all original datasets. We were able to test all possible permutations for all datasets except Or22a.

### InChIKeys as new default odorant identifiers

With this version of DoOR we switched from CAS numbers to InChIs (International Chemical Identifier) and InChIKeys respectively as main chemical identifiers used in DoOR [34]. InChIs are unique chemical identifiers derived from the chemical structure of a compound. InChIs are free to use and the algorithm for generating them is freely available under an open source license http://www.inchi-trust.org/downloads/. Another advantage is that InChIs are human readable and thus their correct use can be verified. As compared to InChIs, CAS numbers are ambiguous: they are assigned to substances rather than compounds. This can result in several CAS numbers for the same chemical. Isopentyl acetate for example, an odorant with a banana like smell for humans, was tested by many studies included in DoOR. Pub-Chem lists 152 synonyms, including “isopentyl acetate”, “isoamyl acetate” and its IUPAC name “3-Methylbutyl acetate”. Among the synonyms are also two different CAS numbers, `123-92-2` as well as `2973250-1` which both map correctly to isopentyl acetate but might have created two separate entries in DoOR. Conversely, the InChI algorithm always produces the standard InChI `InChI=1S/C7H14O2/c1-6(2)4-59-7(3)8/h6H,4-5H2,1-3H3` and the corresponding InChIKey `MLFHJEHSLIIPHL-UHFFFAOYSA-N.` As In-ChIs can be quite long, within the DoOR algorithms we use InChIKeys for all computations. InChIKeys are the 27 character long hashed version of each InChI. While only InChIKeys are used for our DoOR algorithms, we included additional information as a service to the users: we included the name, CID (PubChem Compound Identification) and CAS (Chemical Abstracts Services) identifiers and also added SMILES (simplified molecular-input line-entry system), another structure based chemical identifier.

Redundancy in nomenclature can create duplicates. In fact, even in published sets, we found several cases where the same odorant appeared multiple times in a single dataset, sometimes with different chemical names. Possible explanations could be that the different instances of a chemical were provided from different suppliers. In these cases we merged the entries by calculating the mean responses. Whenever we performed such a merge, we noted that in the `dataset.info` data frame.

### Contribute to DoOR

The source code of the two DoOR packages (`DoOR.data` and `DoOR.functions`) is now available via GitHub (https://github.com/Dahaniel/DoOR.data & https://github.com/Dahaniel/DoOR.functions). This allows to download pre-release versions of DoOR. Everybody (the community) can now contribute feature requests, bug reports and improved code. Package releases will be available via Zenodo (`zenodo.org`) with individual DOIs assigned, thus all `DoOR.data` and `DoOR.functions` versions will be citable. The most recent DoOR version will also be made available *via* the CRAN R-package repository for easy installation from within `R`.

We encourage all users to get access to the full `DoOR.data` and `DoOR.functions`, because they offer several important features, among which direct computer-readable access to all used datasets, routines for back-calculation of consensus responses onto particular datasets, the possibility to add, include or test own datasets and many ways to easily visualize data *via* the DoOR plotting functions. However, we have seen that many users value DoOR for its ease to get immediate responses to quick questions, such as “what is the best ligand for receptor X”, “which are the receptors responding to odorant Y”, or “which glomerulus is innervated by OrZ”. For all of these uses, and graphical displays, we implemented a web interface as a service to the community at http://neuro.uni.kn/DoOR. The interface is largely unchanged with respect to DoOR 1.0, with some changes in the graphical display of the antennal lobe (now based on Grabe *et al.*Grabe2015).

### Broadly and narrowly tuned responding units

How large the set of odors is that a given responding unit is sensitive to can be quantified as its lifetime kurtosis which describes the width of the tuning curve (LTK; Equation 1; Willmore *et al.* Willmore2001). We computed LTK (implemented as the `sparse()` function in `DoOR.functions`) across all DoOR responding units that contained at least 50 odorant responses. We found responding units to be widely distributed across the LTK scale, with many being broadly tuned (low or negative kurtosis; Figure 3a & c) and less that responded only to a few specific ligands (high kurtosis; Figure 3a & b). We found the highest LTK values (most narrowly tuned receptors) for Or82a (LTK = 663.88, n = 180, narrowly tuned to geranyl acetate), ac2A (LTK = 39.12, n = 84, narrowly tuned to putrescine), Or49b (LTK = 35.87, n = 164, narrowly tuned to 2-, 3-, and 4-methylphenol), Gr21a.Gr63a (LTK = 23.57, n = 52, specifically activated by CO_2_) and ab2B (LTK = 20.9, n = 101, tuned to ethyl 3-hydroxybutyrate and cyclohexanol; Figure 3a & b). At the lower end of the scale we found ac3B (LTK = -0.92, n = 98), Or35a (LTK = -0.44, n = 123) (which is expressed in ac3B OSNs and was also measured individually in the empty neuron system), the newly deorphanized Or69a (LTK = -.026, n = 107) and Or85f (LTK = 0.17, n = 114) (Figure 3a & c). The lowest LTK value resulted for ab4B (LTK = -1.53, n = 182), but see below.

**Figure 3:**
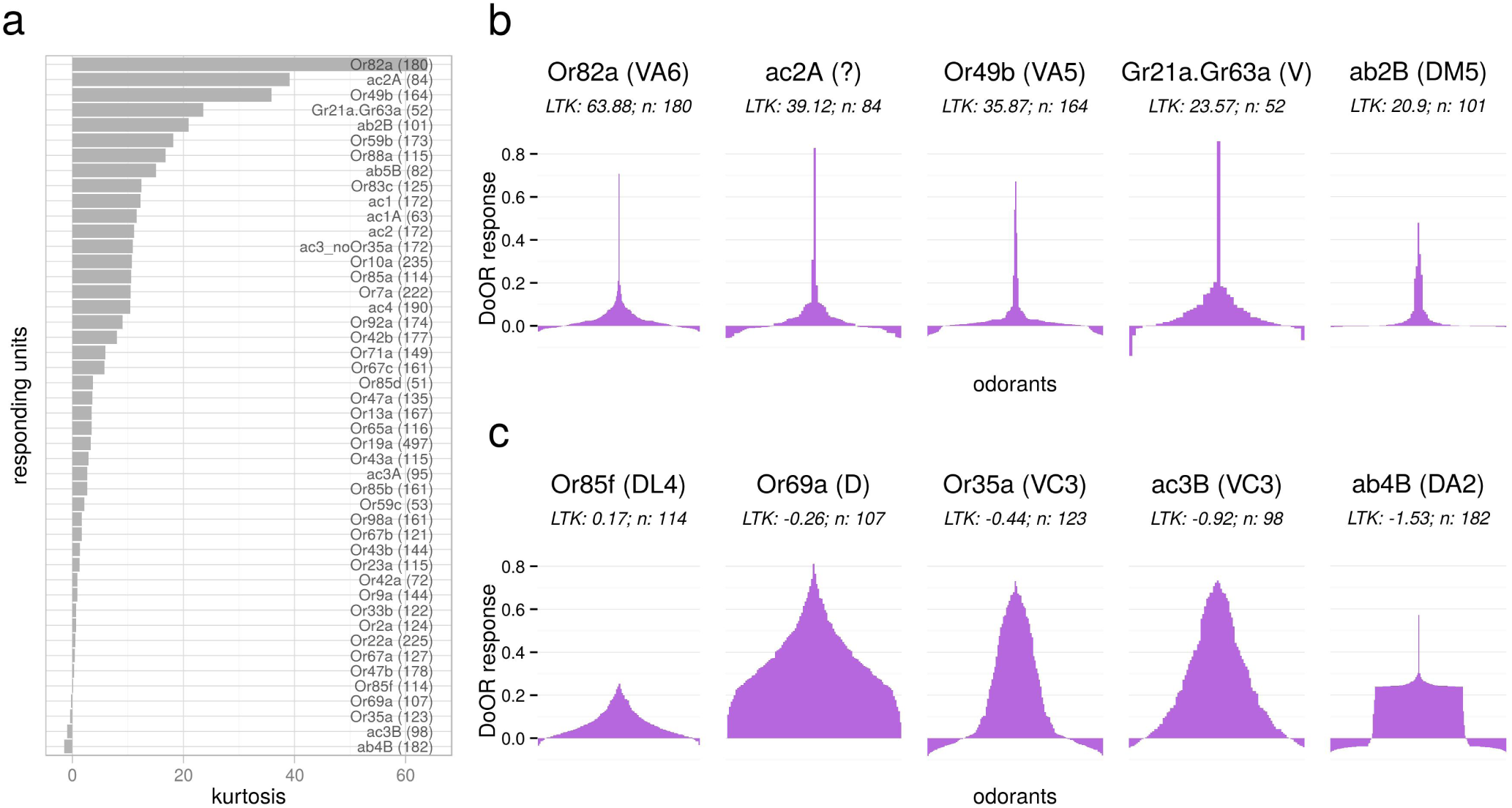
Lifetime kurtosis of DoOR response profiles. **a** Response profiles were ordered according to LTK, higher values indicate sharper odor-response distributions (Equation 1). Only response profiles consisting of at least 50 odorant-responses were considered, the number of odorant responses is given in parentheses. **b** Tuning curves of the five response profiles with the highest LTK. Number of odorants in the dataset is given as *n*. **c** Tuning curves of the five response profiles with the lowest LTK. Corresponding glomerulus names of responding units in **b** and **c** are given in parentheses, names of individual odorants can be seen in the online version of DoOR.

A low LTK value can also indicate that the best ligands for this responding unit is still unknown. For example, Or47b had a low lifetime kurtosis of 0.35 in DoOR 1.0 (where it seemed to be broadly tuned) because at that time only weak responses were known. Dweck *et al*.[27] discovered Or47b to be narrowly tuned to the single compound methyl laurate, for their dataset we calculated the high LTK value of 33.12 (i.e. narrow tuning; Figure S1a). When adding datasets with newly discovered single ligands of narrowly tuned OSNs to DoOR, these responses get systematically underestimated. The reason is in the mathematical model used: merging functions are calculated on the overlapping range of two datasets. If one of the two datasets contains an extremely good ligand, and the other does not, that ligand is, from a statistical point of view, and outlier, and cannot be considered for the merging function. We add these outliers using a linear function with slope 1, added to the fitting function outside of the overlap region. For good ligands, this function creates a systematic underestimation in the consensus set. The situation becomes unfortunately bad when the overlap between two studies (i.e. the odors in common) is low. The Or47b dataset from Dweck *et al*.[27], for example, was excluded from the default merge because it overlapped with all other datasets only by three odorants (the minimum criterion in DoOR is five), and thus the resulting consensus spectrum did not contain the best ligand methyl laurate. Consequently, LTK was low (LTK: 0.25, Figure S1a). When including the Dweck *et al*.[27] dataset by manually adjusting the minimum criterion to three overlapping odors, the resulting LTK value increased to 3.4. This was still lower than the LTK value of 33.12 calculated for the original Dweck [27] dataset (Figure S1b), due to the necessity of mapping to a function with slope 1. We observed a similar effect for ab4B: the LTK value of -1.53 increased to 89.35 when we merged only the studies that included its best ligand geosmin (Figures 3c and S1).

We did not apply any manual selection of source datasets for the pre-computed consensus matrices included in `DoOR.data`, thus Or47b and ab4B have a low LTK in these matrices. We have added a comment to this respect on the homepage. However, we encourage the users to install the DoOR packages and to adjust the merging parameters to their individual needs.

### Odorants activated responding units with differing specificity

We calculated the kurtosis from the perspective of the odorant (population kurtosis, PK). This yields a quantification of how specific or unspecific a given odorant activated the population of responding units (Figure 4). Considering only odorants that were tested with at least half of the responding units present in DoOR (39) we found odorants to be continuously distributed along the PK scale, with some odorants activating small subsets of responding units and many eliciting broad ensemble responses (Figure 4a, b). We found the following odorants on the upper end of the PK scale: CO_2_ activated Gr21a.Gr63a, geranyl acetate activated Or82a, water activated Ir64a.DC4, (1R)-(-)fenchon activated Or85e and methyl salicylate activated Or10a the strongest. The broadest ensemble responses were elicited by 2-heptanone, hexanol, isopentyl acetate, Z3-hexenol and 4-methylphenol (Figure 4a, c).

**Figure 4:**
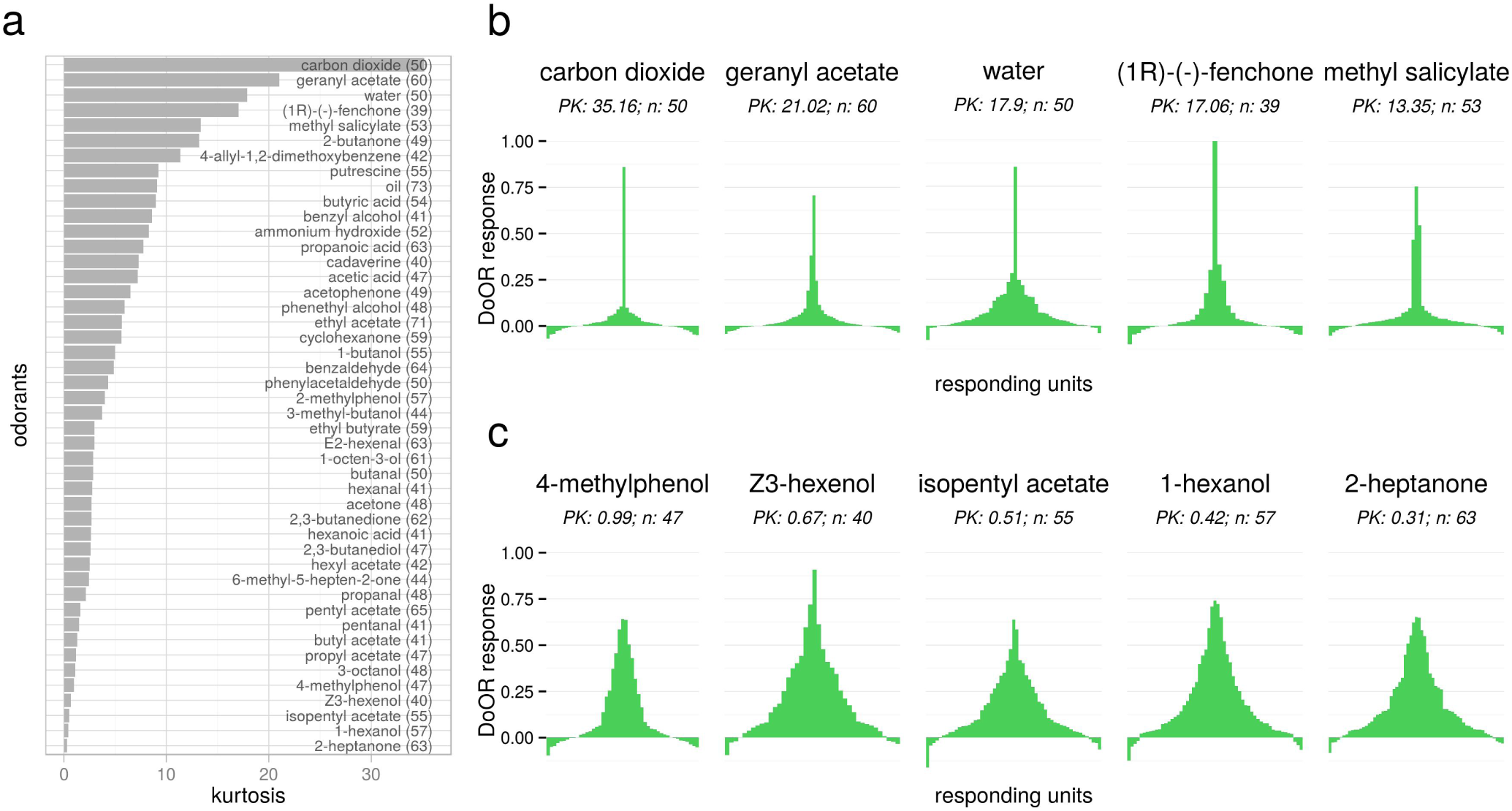
Kurtosis of individual odorants (population kurtosis, PK). **a** Kurtosis was calculated odor-wise, giving a measure of how many responding units were sensitive to a given odorant. **b** Tuning curves of the five odorants with the highest kurtosis. **c** Tuning curves of the five odorants with the lowest kurtosis. *n* indicates the number of responding units known for each odorant. Names of individual responding units can be seen in the online version of DoOR.

### Mapping IRs to their corresponding OSNs

It is difficult to map IR responses to OSNs, because the two available data sources are difficult to disentangle: on one hand single sensillum recordings with spikes that are difficult to sort (with exceptions, in some studies the largest spike amplitude could be separated), on the other hand single IR calcium imaging data, and the difficulty that individual IRs are expressed in several OSN types. See Table 1 for responding units and their relationship to IR/OSN/sensillum/glomerulus. By correlating response profiles from SSR recordings with response profiles from IR calcium data (using the `mapReceptor()` function from DoOR), we have created hypotheses about their respective mappings.

We considered correlations that were significant (p < 0.05) and relevant (correlation coefficient above 0.75). With these criteria we found that glomerulus VC5 (Ir41a) responses correlated with high significance to the DoOR responding units ac2 (summed OSN data; r = 0.8, p-value = 2.3∗10^−6^, n = 24) and ac2A (r = 0.76, p-value =4 ∗ 10^−3^, n = 12). For DP1l the situation is more complex: Two classes of OSNs express Ir75a. Neurons from ac2 sensilla that target glomerulus DP1l express only Ir75a. Neurons from ac3 sensilla that target glomerulus DL2 additionally express Ir75b and c [12]. In our analysis data originating from DP1l correlated with high significance to the ac3A responding unit in DoOR (r = 0.82, p-value = 7.2 ∗ 10^−7^, n = 24), indicating that ac3A neurons expresses Ir75a/b/c and that Ir75a accounts for large parts of the response profile. This supports the notion from Yao *et al.*[35] that the B neuron expresses Or35a. Together the situation appears to be the following: ac2A neurons express Ir75a and target the DL2 glomerulus; ac3A neurons express Ir75a/b/c and target the DL2 glomerulus; ac3B neurons express Or35a (and Ir76b) and target the VC3 glomerulus (Table 1).

While these correlations match published IRsensillum expression patterns [12], we note that Ir64a.DC4 correlated with high significance to ac3A (r = 0.81, p-value = 2 ∗ 10^−6^, n = 24) which would contradict that Ir64a is expressed in sacculus neurons [16, 30]: more experimental data will be needed here.

## Discussion

Animals can code for millions of odors with a limited number of olfactory receptors. Even though estimating the exact capacity of olfactory systems is a matter of fierce debate [36, 37, 38], it is clear that the combinatorial nature of olfaction lies at the basis of this astounding capacity: it is the pattern of activity across receptor neurons that gives the brain the necessary information about the chemicals in its environment. In a combinatorial world (receptor ON or OFF) the theoretical capacity of n receptor types would be 2^n^, with 50 receptors in *Drosophila* that would be 2^50^, corresponding to approx. 10^15^. Since olfactory coding is not binary (i.e. receptors can have all intermediate and also negative activity values), the number might even be higher. On the other hand, the capacity might be smaller, since the code is redundant, and has also to accommodate temporal complexity, concentrations, and mixture analysis, making a prediction of the exact number difficult. Nevertheless, it is apparent that understanding the olfactory code is only possible if the response to a substance is known for all sensory neurons. Given the large number of receptors, and the large number of possible stimuli, this is a daunting task, and no task that a single research group, not even a consortium, could accomplish. Therefore, we have created a technology that allows to merge data from different groups, recorded with different techniques, into a consensus database. The first version has been used by the community for five years now, with great success. We present the second version, with more data and better tools, in this paper.

Every database is a service to the community that needs long-term care and participation. In order to keep its value we need to continuously update and improve data and code and for this rely on the support from the community. For DoOR 2.0 we received considerable help from many colleagues who supplied us with the raw data of their publications, and even with unpublished datasets. Some datasets however could not be obtained (lost data, crashed disks, refusals to reply to emails). DoOR version 2.0 has many improvements over DoOR 1.0. DoOR is now hosted on GitHub so the most recent code and data are always accessible. At the same time releases with individual DOIs will be available *via* Zenodo and CRAN. GitHub also eases bidirectional communication, users can send feature requests, report bugs and hints to missing data *via* the GitHub issue tracker system. InChIKeys are the new unique identifiers used in DoOR, and we added several new functions, e.g. for calculating kurtosis (`sparse()`) or identifying a recorded OSN based on its odorant specificity (`identifySensillum()`). Our goal is to move DoOR increasingly into a common tool used and shaped by the community, rather than just provided by us.

This new version contains additional data for new odorants and additional receptors. DoOR now includes 11 new studies contributing 15 new datasets. 467 new odorants were added, 13 glomeruli and five additional response profiles were deorphanized. In total 2894 responses of new odorant-responding unit combinations were added.

Where are the remaining major lacks in the dataset now? From a neuro-circuitry point of view, the most important lack is in glomerulus VA7m, which has not yet been linked to a receptor. Also, the area innervated by IR receptors in the antennal lobe and their mapping to sensilla and response profiles remains understudied. More data in the next few years will elucidate the missing information. From an olfactory point of view, many odorant responses are still missing (see gray area in Figure 1a). But the number of potentially interesting chemical compounds that may have an odor is virtually infinite, and therefore just adding new compounds will have only limited effect on our understanding of the system. Thus, the major question for the future years will be with respect to those receptors which do not have very strong best responses (e.g. Or2a, Or23a and ab10B). It is conceivable that we are missing the best ligands for these receptors, and that finding a stronger ligand will help us understand the importance of that responding unit within the olfactory code, and for the animal’s ecological niche. These studies will also help us in defining how odors are encoded in the first place, i.e. how the brain extracts information from the activity patterns across receptors. When does the animal interpret activity in a receptor with low kurtosis (e.g. geosmin in ab4B) as that substance being present, when does it interpret activity in the same receptor as a result of another odorant with weaker affinity? The answer most likely is found in the ensemble response across receptors, and can only be found if we know that representation. The value of DoOR lies in the overview of the *Drosophila* olfactome that helps to understand the nature of combinatorial coding. It enables analyses as calculating the kurtosis of ensemble responses elicited by individual odorants or mapping unknown response profiles. It is a resource for modeling, for selecting experimental parameters and can be used for extrapolating new response patterns.

Nevertheless DoOR is - and always will be incomplete, not only in the sense that some odorants will always be missing. Some aspects of olfactory coding are not yet covered, others are impossible to cover in such a consensus approach. Aspects that might be covered in future versions are e.g. odorant concentrations. Merging responses of different odorant concentrations across labs is difficult because it is difficult if not impossible to measure absolute concentration in a controlled way. For one, the absolute odorant concentrations reaching the animal depends on the vapor pressure of a compound. Concentrations are also influenced by how a stimulus is presented. Within studies extremely potent substances are often used in higher dilutions, so that we had to exclude these in some of the DoOR datasets - an unfortunate aspect, because these are, after all, the best ligands. While it is impossible to correctly integrate absolute concentrations of stimuli across studies, it is mathematically possible to integrate concentration-response curves, should they be published more often. In this case, we would add separate datasets per concentration / dilution into DoOR. Similarly saturating and adapted responses created by strong ligands are problematic as they lead to flattening of the odorant response patterns and create distortions in the mapping function. As an example: butyl acetate and ethyl hexanoate elicit equally strong responses from Or22a expressing OSNs when tested at a 1:100 dilution [3] but the OSNs are sensitive to an almost three log steps lower concentration of ethyl hexanoate as compared to butyl acetate (quantified as the dose eliciting the half maximal response; Pelz *et al.*Pelz2006).

Other features that cannot be consensualized in a database include temporal dynamics of odorant responses. However, such information could be implemented in future DoOR releases by enabling to store full time traces as meta links to the individual response points in the matrix, allowing the user to have access to all response time-courses. Similarly, mixture responses or responses to complex odor stimuli are unlikely to enter DoOR due to the many parameters they depend on, but a future version of DoOR might still act as a repository for such information.

There are many other online databases that collect information across studies to provide important tools for neuroscience research. To name a few, FlyBase (http://flybase.org/), and the Virtual Fly Brain (http://www.virtualflybrain.org/) focus on genetic or morphological aspects of *Drosophila*; Sense Lab https://senselab.med.yale.edu/) offers four different olfactory databases offering e.g. odorant activity maps of mouse olfactory bulbs and genetic codes of olfactory receptors of many species; PubChem (https://pubchem.ncbi.nlm.nih.gov/) and ChemSpider (http://www.chemspider.com/) provide detailed information on millions of chemical compounds. All these databases complement each other and the value of each individual database increases with mutual links so that additional information on odorants, genes or morphological structures is only a click away. Currently we link from the individual records of our DoOR web page to FlyBase, the Virtual Fly Brain and PubChem. We plan to increase the linked databases in the future.

DoOR facilitates a variety of analyses on the *Drosophila* olfactome, allowing to ask important questions. As an example, we searched for odorants that elicit narrow activation patterns across responding units (Figure 4c). These odorants are likely of special importance for *Drosophila,* since narrow response patterns are computationally easy to distinguish from patterns elicited by other odorants. One example of such an important odorant for *Drosophila* is CO2, which activates a single type of OSN and innately mediates aversion [39]. Water had the second highest kurtosis value in our analysis. Water is a very important stimulus for every animal (Figure 4c). However the activated Ir64a.DC4 is likely an acid sensor, and whether the water response might be resulting from the pH < 7 of the distilled water remains to be investigated (Ai *et al.*Ai2010; Ana Silbering, personal communication). We also provide a sensillum identification tool, `identifySensillum()`: by inserting odor-response values of a recorded unit, the tool provides plausible responding unit and sensillum candidates. We hope this tool will be helpful for electrophysiologists or other experimenters as a tool for reliably identifying recorded units in physiological experiments.

The increasing number of narrowly tuned OSNs being described in *Drosophila* in the recent years [39, 28, 27] has sparked a debate about whether effective coding of odorants works *via* a set of narrowly tuned labeled line OSNs, or whether the predominant nature of olfactory coding is combinatorial [39, 28, 40, 27, 41]. Drosophil*a* also has many OSNs that are broadly tuned to many different odorants, such as Or35a (in ac3B) or Or69a (in ab9) (Figure 3a & b). Both strategies have advantages. On the one hand being extremely sensitive to ecologically relevant substances is a prerequisite for finding proper food, mating partners or egg-laying sites. On the other hand, broadly tuned OSNs that sample all across the chemical space ensure that animals are not anosmic to new substances and can adapt to new or changing environments. Indeed, the *Drosophila* olfactory system is able to detect and distinguish substances that do not play a role in a fly’s daily life, like explosives, drugs and breast cancer metabolites [29, 42]. It may well be that the same glomerulus that is highly specialized for a particular odorant in a low concentration/high sensitivity mode, participates in an across-glomeruli combinatorial representation in a high concentration/low sensitivity situation [43].

We put this service into the hands of our colleagues and hope to provide a useful tool for the exploration of the olfactory code and for designing future experiments. We strongly rely on your support and feedback and are happy to include new data and process feature requests and bug reports. We also note that the DoOR framework is in no way restricted to *Drosophila* but can be used for different species right away if sufficient odor-response data is available. With the advances in *in-vitro* screenings of human olfactory receptors [44, 45], a human DoOR might soon be possible.

## Material & Methods

### Animals

For new odorant response datasets, all recordings were performed on female *Drosophila melanogaster* expressing the calcium reporter GCaMP 1.3 [46] or GCaMP 3 [47] under the control of the GAL4-UAS expression system. UAS-GCaMP 1.3 flies were provided by Jing Wang, University of California, San Diego, La Jolla, CA; UAS-GCaMP 3.0 flies were provided by Loren L. Looger, Howard Hughes Medical Institute, Janelia Farm Research Campus, Ashburn, Virginia. Stable GAL4-UAS fly lines were of the following genotypes: P[UAS:GCaMP1.3]; P[GAL4:X] (X being one of Or10a, Or13a, Or42b, Or47b, Or67b, Or69a or Or92a), and w;P[Or56a:GAL4]; P[UAS:GCaMP3]attP40.

Flies were kept at 25 ◦C in a 12/12 light/dark cycle at 60-70% RH. Animals were reared on standard medium (100 mL contain: 2.2 g yeast, 11.8 g of sugar beet syrup, 0.9 g of agar, 5.5 g of cornmeal, 1 g of coarse cornmeal and 0.5 mL of propionic acid).

### Odorant preparation

Odorants were purchased from Sigma-Aldrich in the highest purity available. Pure substances were covered with Argon to avoid oxidation. All odorants were applied at 10^−2^ diluted in 5 mL mineral oil (Sigma-Aldrich, Steinheim, Germany). Odorants were prepared in 20 mL head space vials covered with pure nitrogen to avoid oxidation (Sauerstoffwerk Friedrichshafen GmbH, Friedrichshafen, Germany) and immediately sealed with a Teflon septum (Axel Semrau, Germany). A list of all odorants and the measured responses is given in Table S5.

### Calcium imaging

Calcium imaging was performed on two setups which consisted of a fluorescence microscope (BX50WI or BX51WI, Olympus, Tokyo, Japan) equipped with a 50× air lens without cover slip correction (Olympus LM Plan FI 50×/0.5). Images were recorded with a CCD camera (SensiCam, PCO, Kelheim, Germany) with 8 × 8 pixel on-chip binning, which resulted in 80 × 60 pixel sized images. We recorded each stimulus for 20 s at a rate of 4 Hz using TILLvisION (TILL Photonics, Gräfelfing, Germany). A monochromator (Polychrome II or Polychrome V, TILL Photonics, Gräfelfing, Germany) produced excitation light of 470 nm wavelength which was directed onto the antenna *via* a 500 nm low-pass filter and a 495 nm dichroic mirror, emission light was filtered through a 505 nm high-pass emission filter.

### Stimulus application

A computer-controlled gas chromatography auto sampler (PAL, CTC Switzerland) was modified and used for automatic odorant application. A head space of 2 mL was injected in two 1 mL portions at time points 6 s and 9s with an injection speed of 1mLs^−1^ into a continuous flow (60mLmin^−1^) of purified air. The stimulus was directed at the antenna of the animal *via* a Teflon tube (inner diameter 2 mm, length 39.5 cm, with the exit positioned ~ 2 mm from the antenna). Stimuli arrived at the antenna with 750 ms delay due to delays in the autosampler and the flow. Therefor, stimulus onset was determined as 6.75s and 9.75 s.

Four to eight odorants were presented in a row (one block, ISI > 2 min) interspaced by solvent control, room air control and an receptor specific reference odorant. The reference odorants were Or10a: butyl acetate (`DKPFZGUDAPQIHTUHFFFAOYSA-N`), **Or42b**: ethyl propionate (`FKRCODPIKNYEAC-UHFFFAOYSA-N`), **Or47b**: (S)-(+)carvone (`ULDHMXUKGWMISQ-VIFPVBQESA-N`), **Or56a**: 2-hexanol (`QNVRIHYSUZMSGM-UHFFFAOYSA-N`), **Or69a**: isopentanoic acid (`GWYFCOCPABKNJVUHFFFAOYSA-N`). After each injection the auto sampler syringe was flushed with purified air for 30 s. After each block of stimuli, the syringe was washed with hexane or pentane (Merck, Darmstadt, Germany), heated up to 48 ◦C, and rinsed with continuous clean air for 6 min.

### Data analysis: calcium imaging

We analyzed calcium imaging using custom written routines in IDL (ITT VIS, USA) and R[48].

Recorded movies were manually corrected for lateral movement artifacts. Then, an area of interest was defined for the parts of the antenna that showed fluorescence increase upon stimulation. Time traces were averaged across this area. We included all measurements into the analysis as long as animals showed stable responses to the reference odorant.

Relative percentage fluorescence change was calculated as Δ*F*/*F* = ((*F_i_* − *F*_0_)/*F*_0_) × 100 with *F_i_* being the fluorescence at *frame_i_* and *F*_0_ being the mean fluorescence of 5 s before stimulus onset

To correct for the photo-bleaching of the dye, we fitted an exponential decay function of the form *A* ∗ *exp*^−^*^x^*^/^*^B^* + *C* to each response trace using the `nls()` function in R. Because some odorant responses would not reach baseline within measurement time, the decay rate parameter *B* was estimated from the median mineral oil control trace within each animal. We omitted 750 ms at the beginning of the time-trace and 11 s after stimulus presentation. The pre-stimulus part of the recording was weighted 100 fold [49].

Response values were calculated as the mean response during 5 s after stimulus onset subtracted by the mean response during 2.5 s before stimulation.

We corrected for calcium signal decrease over time, likely due to GCaMP bleaching, using a linear regression across reference odorant measurements within each individual animal. The value of this function at each corresponding time point was used to scale responses using the first reference odorant presentation as reference.

Test odorants were measured in n = 3-16 (mean = 8.4) animals. Every individual preparation was used for n = 7-82 (mean = 36.6) different odorants. We averaged all odorant responses across preparations to derive odor-response profiles for the corresponding receptor line.

### Data analysis: merging datasets

The DoOR project builds a consensus database of odor-response profiles from many different laboratories, measured in different measurement units, and comprising incompletely overlapping odorants. Merging these datasets into a consensus odor-response profile was done as described previously [4]. Briefly, the steps were the following: (1) every single dataset was scaled [0,1], (2) two sets with at least 5 common odorants were merged. Merging consisted of fitting 5 different monotonic functions and their inverse (linear, exponential, sigmoid, and two types of asymptotic nonlinear functions, one with an offset and one without) by least-square correlation and then projecting common odors onto the nearest point of the function and unique odors directly onto the function. The fitting function was only used in the range of common data points, and extended beyond this range by a linear function with slope 1. Fit performance of all 10 individual functions was quantified by averaging the orthogonal distances between the original position of the common points and their final position on the fitted function (“mean distance”, MD). This resulted into a reduction of the number of datasets by 1, since two sets were merged. (3) step 2 was repeated, until only one set remained.

With this procedure, the sequence of merging is relevant. Therefore, wherever computationally feasible, we ran all possible permutations for merging. We evaluated the result of each permutation by calculating the average MD of the merged dataset against every single dataset. The permutation that gave the least overall MD was chosen, and the resulting consensus dataset appears in DoOR. In the other cases, all pairwise merges between datasets were calculated and the pair with the lowest MD was merged. This step was then repeated until all datasets were merged. Datasets that had less than five overlapping odorants with other datasets, and datasets that did not yield a MD below a defined thresh-old (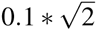, which corresponds to 10 % of the maximum possible distance within the square [0,1] response space) were excluded from the merging process.

In the resulting consensus dataset the responses within each receptor were scaled [0,1]. Using information from studies that recorded more than one receptor we next performed a global normalization that rescaled response ranges of receptors relative to each other. This final consensus dataset is scaled [0,1] across all receptors but as a result of global normalization the individual responding unit will not fill the full range. Many OSNs tend to fire action potentials even in the absence of odorant stimuli. These background or spontaneous firing rates (SFR) are often reported in electrophysiological studies. We treated SFR as a normal odorant. For all studies that subtracted but did not report SFR or for calcium imaging studies, we set SFR to 0. To regain negative response values (inhibitory odorants) we then subtracted the individual SFR values from all other response values (e.g. using the `resetSFR()` function).

### Data analysis: analysis of response properties

We quantified the widths of OSN tuning curves by calculating the lifetime kurtosis (LTK, sometimes referred to as lifetime sparseness) as follows:

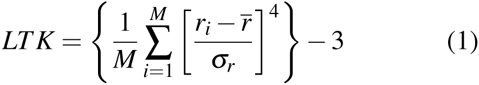

with *M* being the number of stimuli, *r_i_* the response elicited by stimulus *i* and 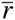 and *σ_r_* the mean and the standard deviation of the responses [50]. High values indicate narrow tuning curves, a kurtosis of 0 corresponds to the Gaussian distribution.

Similarly, we quantified population kurtosis (PK) using the same formula, but across response units, for each odorant. This statistic is sometimes referred to as population sparseness.

### DoOR code

All DoOR code was written in R. The code is available on GitHub (https://github.com/Dahaniel/DoOR.functions), and every modification is documented. Chemical identifiers received from colleagues in different formats were translated into InChI and InChIKeys *via* the cactus service (http://cactus.nci.nih.gov/) using the webchem package (https://cran.rstudio.com/web/packages/webchem/). When no hit could be found, substances were looked up manually at Pub-Chem (https://pubchem.ncbi.nlm.nih.gov/), or ChemSpider (http://www.chemspider.com/).

The graphical display of odor-responses in the antennal lobe was changed, and is now based on a 3D-Atlas by Grabe *et al.* [17].

Plotting was performed using the ggplot2 package [51].

## Acknowledgments

We are extremely thankful to all colleagues that supplied us with the original data from their publications. Ana Silbering contributed unpublished data which we highly appreciate. We thank Veith Grabe for contributing an antennal lobe map optimized for mapping the DoOR odorant activity. We thank Jennifer Ignatious-Raja, Michael Thoma, Shouwen Ma and Tom Laudes for help with the physiological measurements. We thank Eduard Szöcs from rOpenSci for pointing us to his webchem project that we used for retrieving and translating chemical identifiers automatically which likely saved hundreds of working hours. We acknowledge support by the German Research Foundation, DFG.

## Contributions

D.M. performed experiments, analyzed data and wrote the code. D.M. and C.G.G. designed research and wrote the manuscript.

